# Actionable, Motivating, and Not Beyond Question: Student Insights On AI-Enabled Feedback and Instructor AI Use

**DOI:** 10.64898/2026.11.13.766505

**Authors:** Michele Weston, Tammy Long

## Abstract

Scientific writing helps students develop and communicate their understanding through practices such as constructing explanations and arguing from evidence, but its complexity often requires individualized feedback and guidance. This makes scientific writing a worthwhile context in which to study the effects of feedback augmentation with generative AI (genAI). Although emerging work demonstrates that genAI can support feedback generation, considerably less is known about how students experience and respond to AI-enabled feedback. At the same time, feedback research has identified characteristics students view as supportive, yet it remains unclear whether these insights apply to AI-enabled feedback. Accordingly, this qualitative study examines biology students’ experiences and perceptions of feedback developed by their instructor with assistance from a custom genAI tool in an authentic learning context. Analysis of student team interviews developed three themes: students recognized characteristics of effective feedback, positive emotions encouraged intention to revise, and students accepted but critically evaluated instructor AI use. Together, the themes suggest that AI-enabled feedback supported revision through both its content and its influence on students’ academic emotions, which contributed to more favorable views of instructor genAI use. However, students’ evaluations also depended on whether AI use aligned with their expectations and on the credibility of both the instructor and the genAI tool, for which instructor oversight was an important consideration. Collectively, these findings highlight the importance of a learner-centered approach to genAI implementation that accounts for students’ needs and expectations and promotes engagement with feedback. Together, these findings suggest that students’ positive perceptions of AI-enabled feedback were rooted less in the use of AI itself and more in the extent to which the feedback supported revision and improvement.

## 1. Introduction

Modern perspectives on undergraduate biology learning and assessment increasingly emphasize three-dimensional learning (Laverty et al., 2016; Pellegrino et al., 2014). This approach is grounded in frameworks that propose students develop a deeper understanding of scientific ideas by engaging in the practices of inquiry and discourse through which those ideas are developed and adapted (American Association for the Advancement of Science, 2011; National Research Council, 2012). At the same time, scientific knowledge becomes meaningful and useful when students apply it through those practices to investigate and explain the natural world (American Association for the Advancement of Science, 2011). Scientific writing can support three-dimensional learning by providing opportunities for students to develop and communicate their understanding while engaging in scientific practices such as constructing explanations and arguing from evidence. However, scientific writing is a multifaceted and challenging task. Students must navigate disciplinary conventions while clarifying and communicating scientific ideas for a specific purpose and audience (Rivard, 1994).

Given the complexity of scientific writing, students require personalized feedback and guidance from instructors (Jerde & Taper, 2004; Yeh, 2010). Although students have clear views about what makes feedback effective, these preferences do not always align with the feedback they receive, reducing their motivation to use that feedback (Ferguson, 2011; Jørgensen et al., 2023; Winstone et al., 2017). This means that students often continue to receive feedback that does not support their learning and efforts to improve (Carless, 2006; Derham et al., 2022). A contributing factor may be the demands that feedback places on instructors. Providing detailed, personalized guidance is a time-intensive process, particularly in courses with large enrollments (Hanmore et al., 2024). Unfortunately, biology instructors may choose to omit scientific writing from curricula when they do not have the time and resources to provide quality feedback (Jerde & Taper, 2004).

Given the many challenges involved in developing longer pieces of scientific writing, and in the feedback process, it is a worthwhile context in which to study the effects of feedback augmentation with generative AI (genAI). Instructors are increasingly using genAI to help develop student-facing feedback on scientific writing, such as lab reports (e.g., Mills et al., 2025; Wan & Chen, 2024), research proposals (Weston & Long, 2026) and shorter constructed-response tasks (e.g., Chu et al., 2026; Cooper & Klymkowsky, 2024; Pellegrino, 2024). Although this emerging work demonstrates that genAI can support feedback generation, considerably less is known about how students experience and respond to AI-enabled feedback. Recent studies have examined students’ perceptions of AI feedback within a variety of implementations, and findings have been mixed. In some contexts, students find AI feedback on scientific writing to be useful, actionable, and clearly written (Mills et al., 2025). In other cases, however, students perceive the feedback as less effective. For example, Lim and Go (2025) reported that students found AI-generated feedback on essays to be overly broad, generic, and vague. One possible explanation for these differences is that Lim and Go relied on a standard ChatGPT prompt without customization, resulting in feedback that consisted of generalized comments resembling grading criteria. In contrast, Mills et al. (Mills et al., 2025) used a customized version of ChatGPT equipped with specialized instructions and a curated knowledge base, enabling it to generate detailed and consistent feedback.

This gap in understanding students’ perceptions of AI feedback, and their experiences using the feedback to revise their work, make it unclear how genAI influences the feedback process. An exploratory study of students’ perspectives within an authentic learning context is particularly valuable for generating hypotheses and tentative explanations of how students are affected by AI integrations in the feedback process. Accordingly, this exploratory qualitative study examines biology students’ experiences and perceptions of feedback developed by their instructor with the aid of a custom genAI tool, which we refer to as AI-enabled feedback. Specifically, we asked:

When biology undergraduates are provided AI-enabled feedback on their scientific writing:

a. How do they experience and perceive the feedback itself?
b. How do they perceive instructor use of genAI for feedback?

### 1.1 Learner-Centered Feedback Process

Research on feedback has shifted considerably over time. Earlier conceptualizations defined feedback as information provided to students about their performance, particularly with the intention of closing the gap between their current and the desired performance levels (Hattie & Timperley, 2007). Within this view, feedback is treated largely as something instructors deliver to learners, leading researchers to focus on characteristics of the feedback itself, such as its content, timing, and mode of delivery.

More recent conceptualizations have challenged this transmission-oriented perspective by emphasizing the role of learners in making feedback effective. Rather than viewing feedback as information that is given to students, learner-centered approaches define feedback as a process through which students also make sense of information about their performance and use it to improve their future learning (Boud & Molloy, 2013; Henderson et al., 2019). From this perspective, the value of feedback depends not only on what is communicated but also on how learners interpret and act on that information. Consequently, feedback research has increasingly examined students’ experiences with feedback, including how they understand it, respond to it, and incorporate comments into subsequent work (e.g., Dawson et al., 2019; Jonsson, 2013; Li & De Luca, 2014). Consistent with this perspective, we conceptualize feedback as a learner-centered process in which outcomes depend on students’ experiences, engagement with, and use of feedback.

### 1.2 Student Perceptions of Effective Feedback

Student perceptions of the characteristics that make feedback effective are important considerations within a learner-centered feedback framework. These perceptions are shaped, in part, by students’ understandings of the purpose of feedback. Students often view the primary purpose of feedback as helping them improve their work (Dawson et al., 2019; Jonsson & Panadero, 2018; Li & De Luca, 2014; Still & Koerber, 2010). Consistent with this perspective, prior research has shown that students find feedback useful when it explains how to make improvements, particularly when they have opportunities to revise their work or apply comments to future assignments (Dawson et al., 2019; Higgins et al., 2002; Jonsson & Panadero, 2018; Li & De Luca, 2014; Lipnevich & Smith, 2009; Lizzio & Wilson, 2008; Weaver, 2006).

Research suggests that students find feedback most useful when it is comprehensive, actionable, and aligned with assignment criteria (Carless, 2006; Dawson et al., 2019; Lipnevich & Smith, 2009; Lizzio & Wilson, 2008; Weaver, 2006). Comprehensive feedback provides sufficient context for students to understand both the nature of a problem and what needs to change, reducing ambiguity. For example, a student in Carless’s (2006) study criticized the comment, “you need to elaborate more,” explaining that they “had no idea what more I could say.” In addition, students find feedback actionable when it provides enough information, often through examples or hints, to help them improve their work (Dawson et al., 2019; Lipnevich & Smith, 2009; Weaver, 2006). Students also report finding feedback more productive when it is aligned with assignment criteria (Dawson et al., 2019; Lizzio & Wilson, 2008; Weaver, 2006), as this alignment clarifies how addressing the feedback can reduce the gap between their current and expected performance. In contrast, feedback that is poorly aligned with assignment criteria may leave students uncertain about priorities for improvement.

Students also tend to perceive feedback as more effective when it provides sufficient detail (Dawson et al., 2019; Ferguson, 2011; Van der Kleij & Lipnevich, 2021; Weaver, 2006), with some students identifying detail as its most important characteristic (Lipnevich & Smith, 2009). Detailed feedback is considered effective because it reduces confusion and helps students understand how to improve their work. In contrast, students often view one-word comments, unexplained feedback, and markings such as ticks, circles, underlining, and other symbols as unhelpful (Ferguson, 2011). Students generally prefer either a written summary at the end of an assignment or brief comments embedded throughout their work. Embedded comments can be particularly useful because they help students connect feedback to specific parts of their writing (Ferguson, 2011).

Beyond providing actionable, detailed comments, students identify several other characteristics that contribute to effective feedback. They prefer feedback that is personalized, timely, and fair. Personalized feedback is more actionable because it offers recommendations tailored to a specific piece of writing rather than simply restating assignment criteria or providing generic comments to an entire class (Ferguson, 2011; Lipnevich & Smith, 2009). Students also view personalization as evidence that an instructor cares about their learning and has invested time in reading their work and providing quality feedback (Lipnevich & Smith, 2009; Lizzio & Wilson, 2008). Timeliness is another important consideration (Ferguson, 2011; Van der Kleij & Lipnevich, 2021). Students recognize that feedback is most useful when it is received before the next assignment and while there is still an opportunity to apply it (Ferguson, 2011). They may also prefer feedback shortly after submission, with participants in Ferguson’s (2011) study suggesting a turnaround time of two to three weeks. Students also desire that feedback is fair, although what constitutes fairness is not always clearly defined (Li & De Luca, 2014). In Carless’ (2006) study, students described fair feedback as unbiased, which they described as unaffected by the instructors’ perceptions of students’ overall performance in the class, and free from favoritism. They also viewed it as unfair when some peers received more support than others. In addition, students in Lizzio and Wilson’s (2008) study associated fairness with instructors providing justification for their comments.

The characteristics above help make feedback supportive of improvement, but students also commonly desire feedback that is mindful of affect (Coleman et al., 1987; Ferguson, 2011; Lizzio & Wilson, 2008; Van der Kleij & Lipnevich, 2021; Weaver, 2006). This topic is explored further in the section “Feedback and Academic Emotions”, but here we focus on students’ perceptions of how the positive and negative valence of feedback shapes their emotional responses and overall experiences. Positive comments can provide socioemotional support that helps students manage the affective dimension of receiving feedback (Hattie & Timperley, 2007; Lipnevich & Smith, 2009; Lizzio & Wilson, 2008). For example, most students in Weaver’s (2006) study agreed that positive feedback boosted their confidence and self-esteem. Students typically regarded comments as positive when they acknowledged achievements, recognized effort, conveyed criticism in a considerate tone, or fostered hope for improvement.

Students’ perceptions of negative feedback are nuanced and often depend on the balance between critical and positive comments (Coleman et al., 1987; Weaver, 2006). Weaver (2006) found that students’ responses to negative feedback were generally less severe than expected; however, feedback with mostly negative comments was viewed less favorably and was often described as demoralizing. Students also reported that feedback frequently focuses only on weaknesses while overlooking strengths, and many suggested that feedback could be improved by including more positive comments. Taken together, these findings suggest that students perceive feedback more favorably that balances constructive criticism with acknowledgment of their effort and accomplishments.

Student perceptions of the feedback sender can also influence their perceptions of feedback effectiveness, although this perspective remains understudied (Orsmond et al., 2005; Poulos & Mahony, 2008; Telio et al., 2016). Research suggests that when students perceive a feedback sender as credible, they are more likely to engage with the feedback (Telio et al., 2016; Watling et al., 2012). Similarly, Ruwe and Mayweg-Paus (2023) compared students’ perceptions of the trustworthiness of feedback generated by an AI system, an instructor, or a peer. The authors found that students viewed the AI system, either acting independently or supporting the instructor, as the most trustworthy feedback provider because it was perceived as having greater expertise. More research is needed to better understand how students’ perceptions of the feedback sender influence their experiences with and use of feedback, particularly in the context of AI-enabled feedback.

### 1.3 Feedback and Academic Emotions

Beyond cognitive processes, such as understanding and interpreting feedback, students also respond to feedback emotionally, which can shape its effectiveness. For example, feelings of confidence and competence can support students’ willingness to re-engage with a task following feedback (Ferguson, 2011; Hattie & Timperley, 2007). In contrast, feelings such as helplessness or resignation may limit students’ receptiveness to feedback and their willingness to invest effort in improving (Brookhart, 2017; Ferguson, 2011; Kernis et al., 1989; Linnenbrink, 2007).

For the present study, we adopt Pekrun and Linnenbrink-Garcia’s (2012) definition of academic emotions as emotions that arise in relation to learning activities or their outcomes. Outcome-related emotions are especially relevant in the context of feedback because feedback focuses on students’ performance on a completed learning task. Consequently, students’ emotional responses to feedback can be understood as both a component of and an influence on their broader reaction to that performance outcome (Pekrun, 2006).

According to Control-Value Theory, academic emotions arise from students’ personal appraisals of control and value (Pekrun, 2006). These appraisals relate to students’ perceived control over their success on learning tasks, and the values they place on the tasks and outcomes, respectively. Different combinations of control and value appraisals contribute to different emotional responses, helping explain why students may react differently to similar learning experiences. In the context of feedback, students’ emotions may vary depending on the extent to which they believe they can improve their work and view improvement as worthwhile or important (Pekrun, 2006).

Emotions can affect learning in different ways, and researchers often conceptualize them along two dimensions: valence (positive versus negative) and activation (activating versus deactivating; Pekrun, 2006; Pekrun & Linnenbrink-Garcia, 2012). Positive activating emotions motivate students to direct effort toward the learning task, whereas negative deactivating emotions reduce students’ motivation to do so. Conversely, the effects of positive deactivating emotions and negative activating emotions on learning is more complex. For example, anxiety is generally considered a negative emotion, yet it may either motivate increased effort or contribute to avoidance, depending on how it is experienced in a particular context. Considering both dimensions helps explain why emotions with the same valence can have different effects on learning. Because emotions can shape students’ willingness to devote effort to learning tasks, they are theorized to influence motivation and, ultimately, achievement (Pekrun, 2006; Pekrun & Linnenbrink-Garcia, 2012).

This relationship between emotions and motivation is particularly relevant in the context of feedback and revision. Students’ achievement on subsequent drafts may be influenced by how their emotional responses affect their willingness to engage with feedback and revise their work (Pekrun, 2006). Because the present study relies on interviews rather than revision artifacts, we focus on students’ motivation to revise. In a prior analysis of the revisions produced by the same students participating in this study, we found that students frequently acted on the AI-enabled feedback and made substantive changes to their writing (Weston & Long, 2026). However, revision products alone could not reveal how students experienced the feedback or how their experiences may have influenced their motivation to revise. The present study builds on these findings using interviews to examine students’ experiences with AI-enabled feedback, including their reflections on the feedback itself and using it to revise their scientific writing.

### 1.4 Generative AI Use in Feedback Development

Instructor feedback plays a crucial role in supporting students’ scientific writing, and recent advances in genAI have created new opportunities to assist instructors in producing such feedback (Mills et al., 2025; Wan & Chen, 2024). GenAI systems are artificial intelligence tools that generate novel outputs, including written text, in response to user prompts (Stryker & Scapicchio, 2024). Several features make genAI appealing for feedback generation. Unlike many traditional machine-learning approaches, genAI systems can be directed through natural-language instructions and do not require large task-specific datasets to develop feedback. Because instructors can interact with these systems through prompting and customization rather than model training, genAI makes AI-supported feedback more accessible to instructors and may support its broader adoption.

## 2. Methods

This qualitative study is grounded in a contextualist epistemology, which emphasizes the context-dependent and multilayered nature of language and meaning (Madill et al., 2000). This paradigm recognizes that researchers cannot be separated from their values and background and that these inevitably shape the research process (Cruz, 2015). Together, these assumptions suggest that the same evidence may reasonably support multiple interpretations (Cohen et al., 2018). This epistemological perspective has implications for our study design. Specifically, we conducted this study in a naturalistic setting to preserve the contextual complexity of participants’ experiences (Hammersley & Atkinson, 1995), rather than in a controlled experimental setting that requires standardization and researcher manipulation.

We used a multiple case study design to investigate students’ experiences and perceptions of AI-enabled feedback. Multiple case studies emphasize an in-depth analysis of a small number of cases, where the cases are emphasized over specific variables (Ragin & Becker, 1992). As such, case studies are ideal for presenting insights that inform our understanding of the nuance of the experiences of student teams with AI-enabled feedback and their perceptions of instructor AI use (Merriam, 1998; Miles & Huberman, 1994). The study includes two student team cases, each consisting of a team of three students.

### 2.1 Trustworthiness and Reflexivity

Consistent with quality criteria for qualitative research (Braun & Clarke, 2006), we sought to establish rigor through transparency, systematic analysis, and reflexivity. Transparency was supported through detailed reporting of the study procedures and analytic decisions, while systematicity was supported through a structured and documented analytic process described below. To support reflexivity, defined here as an examination of how one’s experiences, assumptions, and context may influence the research process (Cruz, 2015), the first M.W. maintained a reflexivity journal throughout the study. The reflexivity statements presented below offer insight into the researchers’ interpretive frames that shaped the study’s analytic process.

#### 2.1.1 Research Team Reflexivity

Our research team brought differing experiences with K-12 and undergraduate science education, which were used to challenge interpretations and refine themes. We entered this study with a broad interest in the use of generative AI to support formative feedback on scientific writing, including its effect on student experiences and perceptions. We recognized that these interests could sensitize us to positive accounts of AI-enabled feedback. To examine this influence, M.W. used analytic memoing and reflexive journaling, revisited disconfirming cases, and discussed alternative interpretations during theme development.

The backgrounds of individual team members also shaped the perspectives brought to this work. At the time of the study, M.W. was a PhD student with prior experience as a high school science teacher. Her earlier research focused on automated analysis of constructed-response questions in biology using text analysis and supervised machine learning techniques. These professional experiences influenced her perspective on formative feedback processes and the role of AI in education. As a graduate student, she regularly used LLM-powered tools such as ChatGPT and Copilot to support her academic work and learning. This firsthand use of genAI provided experiential insight into the student perspective on AI in education.

T.L. was a tenured professor and biology education researcher with 25 years of teaching experience in undergraduate biology. Her formal training was in plant ecology but post-Ph.D., her research focus shifted toward education and learning in the biological sciences. She has minimal experience with AI tools and at the time of the study, had not used them to support either her teaching or research.

### 2.2 Study Context

We conducted the study within a STEM-focused residential college at a large Midwestern research university. Data were collected during the laboratory component of a combined lecture–laboratory introductory organismal biology course in Spring of 2025. The course was comprised mostly of second-year students. The course curriculum was designed to explore systemic inequities and human impacts on ecology and health.

Students were grouped into teams that stayed the same and worked together through the entire course. Teams were assigned using a combination of student requests and recommendations from the CATME Team-Maker system (*CATME Project*, n.d.; Layton et al., 2010), which was configured to incorporate the factors including schedule, demographics, leadership, and writing style. Teams spent the second half of the course developing independent research projects investigating the underlying ecological cause of a health disparity of their choice. The research projects accounted for 56% of students’ grade in the lab component of the course, and the lab component accounted for 27% of their overall course grade. As a part of the research project, teams wrote the introduction section of a research proposal as a group assignment. Therefore, in this report we will refer to the piece of writing as the “proposal introduction”. Students were instructed to follow a rubric that specified the content of four required paragraphs, each focusing on a different scale of human impact. Descriptions of the paragraphs are shown in Table 1.

**Table 1.** Content of the paragraphs of the proposal introduction.

| Paragraph Name | Description of Content |
|---|---|
| Macro | This was the first paragraph of the proposal introduction. Students were to describe the problem at the macro (historical and societal) scale. They were to explain how historical practices (e.g., redlining) result in current racial disparities in health outcomes and local ecosystems. |
| Meso Part 1 | This was the second paragraph of the proposal introduction. Students were to describe the problem at the meso (community) scale. They needed to explain the impacts of specific policies or practices (e.g., urban renewal, highway construction) on local resources, community health, and environmental conditions, focusing on neighborhood structures and resources. |
| Meso Part 2 | This was the third paragraph of the proposal introduction. Students were to describe one or more community responses to inequity that are relevant to the experimental question and variables. |
| Micro | This was the fourth paragraph of the proposal introduction. Students were to describe the problem at the micro (individual) scale, and explain how systemic inequities affect personal health outcomes and ecological interactions, highlighting specific cases and personal experiences (e.g., personal stories of residents in affected areas). |

### 2.3 Participants

All students enrolled in the laboratory course were invited to participate in the research study. Two complete teams consented to participate and are considered team cases, each comprised of three students. Table 2 presents demographic and academic information for the student participants. Gender and race(s) and ethnicities were self-reported. Final course grade and cumulative GPA were obtained by the Registrar’s office with consent from the participants.

**Table 2.**
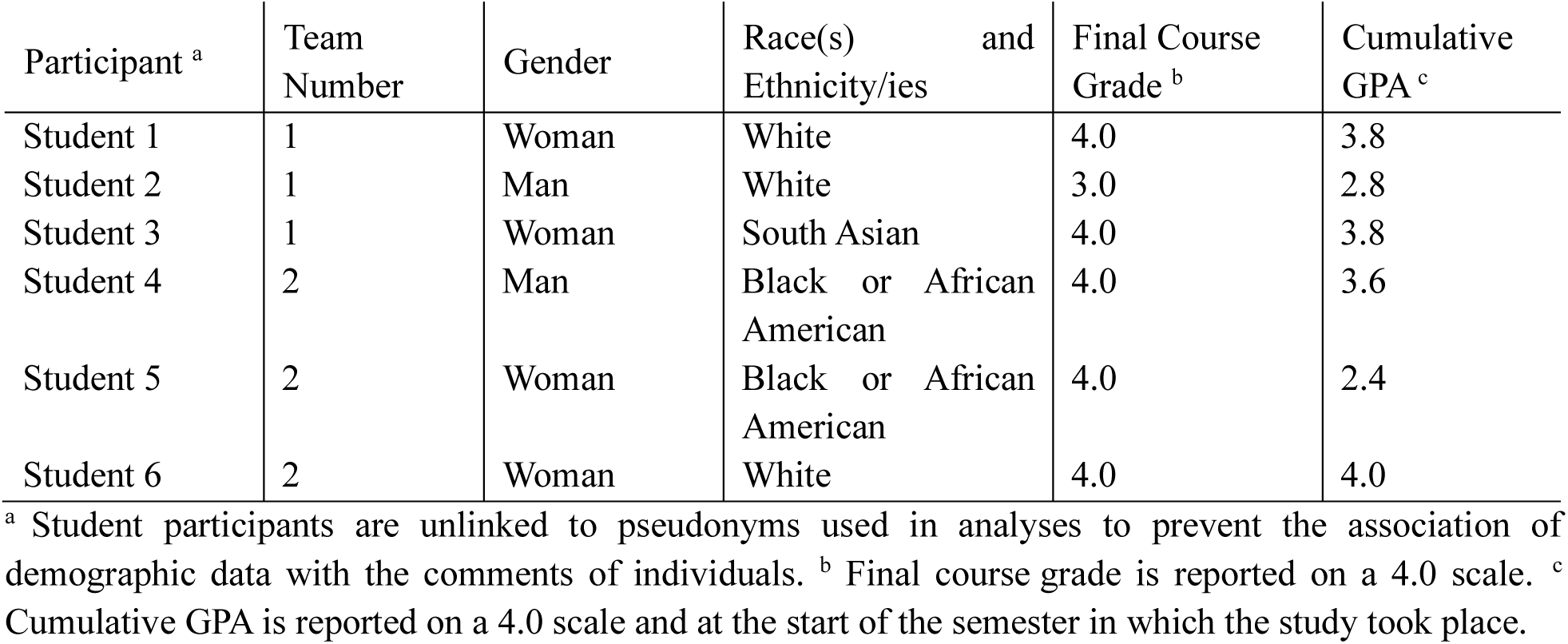
Student participant demographics.

| Participant<sup>a</sup> | Team Number | Gender | Race(s) and Ethnicity/ies | Final Course Grade<sup>b</sup> | Cumulative GPA<sup>c</sup> |
|---|---|---|---|---|---|
| Student 1 | 1 | Woman | White | 4.0 | 3.8 |
| Student 2 | 1 | Man | White | 3.0 | 2.8 |
| Student 3 | 1 | Woman | South Asian | 4.0 | 3.8 |
| Student 4 | 2 | Man | Black or African American | 4.0 | 3.6 |
| Student 5 | 2 | Woman | Black or African American | 4.0 | 2.4 |
| Student 6 | 2 | Woman | White | 4.0 | 4.0 |

<sup>a</sup> Student participants are unlinked to pseudonyms used in analyses to prevent the association of demographic data with the comments of individuals. <sup>b</sup> Final course grade is reported on a 4.0 scale. <sup>c</sup> Cumulative GPA is reported on a 4.0 scale and at the start of the semester in which the study took place.

The instructor was a Teaching Assistant Professor and had three semesters of prior teaching experience. She had not used genAI prior to this study.

#### 2.3.1 Case Description of Student Team 1

Team 1 examined the research question, “In previously redlined communities, what is the relationship between tree density and median income, asthma frequency, and UV exposure?” in the context of a local community. Their proposal introduction identified the construction of a nearby highway as a historically connected event that continues to shape environmental and health outcomes. The team also highlighted a local garden program as a community-based response that may help address harms associated with both redlining and highway development.

#### 2.3.2 Case Description of Student Team 2

Team 2 investigated the question, “How has redlining affected community garden quality and prevalence in the greater [city] area and how is it tied to obesity?” Their proposal emphasized redlining and characterized obesity as a long-term health outcome linked to it. To examine this relationship, they discussed how poor soil quality and limited access to grocery stores in historically redlined communities may contribute to higher obesity rates. The team also described nearby community gardens that donate produce to a local food bank as a response initiative designed to reduce environmental and health disparities.

### 2.4 Procedures

Student teams began the assignment by drafting the proposal introduction and submitting it to their instructor. The instructor then used a genAI tool (detailed in section 2.6) to develop customized feedback aligned with the assignment rubric, reviewed the feedback for accuracy and quality, and edited it before providing it to students. The instructor had the option of using follow-up prompts to refine the genAI tool output but opted to only make a personal edit. The instructor made one edit to Team 1’s feedback and did not edit Team 2’s feedback. Students then revised their drafts in response to the feedback and submitted their final drafts. At the start of the course, students were informed of their instructor’s use of genAI to assist with developing feedback. However, it came to light during student team interviews that both teams had forgotten this discussion by the end of the semester. Consequently, students were not aware their feedback was developed with the assistance of genAI until after they made revisions, and at the start of the student team interviews with the researcher.

#### 2.4.1 Student Interviews

We conducted semi-structured small group interviews with each student team to explore their experiences and perceptions of the AI-enabled feedback. The interview protocol consisted of three parts, including questions about: 1) students’ prior experience using genAI and their initial perceptions of the instructor’s use of genAI for feedback; 2) students’ views of the AI-enabled feedback itself; and 3) students’ thoughts, feelings, and concerns about instructor use of AI following this project. The interview protocol is provided in the Supplemental Materials. Interviews were transcribed using Zoom and subsequently edited for accuracy by research assistants.

### 2.5 Data Analysis

We analyzed interview transcripts using reflexive thematic analysis (Braun & Clarke, 2022). This approach offered an inductive experiential analysis, with both descriptive and interpretive accounts of the data. However, reflexive thematic analysis’ analytical flexibility also allowed for a shift to a more deductive orientation when necessary, which we turned to during theme development as described below.

For our analysis we followed the procedures for reflexive thematic analysis outlined by Braun and Clarke (2022). Accordingly, the analysis was iterative and consisted of the following steps. First, M.W. read the interview transcripts twice and took familiarization notes on her initial perceptions of the data. Next, she systematically coded the transcripts in MaxQDA (VERBI Software, 2024), inductively applying code labels that summarize the idea being communicated, and across segments containing similar ideas. After coding, M.W. began using these coded patterns to develop themes that depicted her interpretation of shared meanings across the data. To promote the quality of the analysis, both authors met periodically during theme development to discuss varying interpretations and bring broader perspectives to bear on the data. Although theme development was largely inductive, we used academic emotion theory (Pekrun, 2006; Pekrun & Linnenbrink-Garcia, 2012) as an analytic lens during the development of Theme 2, reflecting a more deductive approach. This was because students frequently discussed emotional reactions to the AI-enabled feedback, so we applied the theory to help interpret how those emotions related to students’ experiences with feedback and revision.

### 2.6 Transparency, Ethics, and Pedagogy Considerations of Using AI

We report the present study following the guidelines of the Transparency-Ethics-Pedagogy-AIED (TEP-AIED) model, which promotes transparent, ethical, and pedagogically grounded applications of AI (Hwang et al., 2026). To support transparency, we document the genAI tool used, including its provider, version, and the instructions and configurations (see below and in the Supplemental Materials). From an ethical perspective, we report ethical approval and informed consent procedures, describe measures taken to anonymize and protect participant data, and consider potential risks associated with AI use, including hallucinations. Pedagogically, we outline the intended learning objectives, the pedagogical framework, and instructional supports informing the assignment and custom AI instructions, and the specific role of genAI within the feedback process. These considerations guided both the design of the implementation and the interpretation of findings related to students’ perceptions and experiences of the feedback.

We utilized a GPT, a specialized version of ChatGPT developed to perform a specific task (*Introducing GPTs*, 2024), which utilized the large language model GPT-4o (OpenAI, 2025). Although the GPT resembled the standard ChatGPT interface, where the instructor interacted with a chatbot through a chat window, it operated under predefined instructions and incorporated a knowledge base specific to the research proposal assignment. GPTs are well-suited for developing customized feedback because they can follow a prescribed analysis protocol, reference internal documents such as rubrics, access topic-specific information, justify their outputs, and respond to instructor queries. In addition to providing written consent to participate in the study, participants provided written consent for the use of genAI in analyzing their work after being informed of the procedures and genAI’s role in the project. To protect student privacy, we anonymized all submissions prior to analysis and we excluded proposals from OpenAI’s training dataset.

#### 2.6.1 Human-Centered Approach to AI-Enabled Feedback

As genAI becomes increasingly integrated into educational practice, questions of quality and responsible implementation have become essential. One approach to addressing these concerns is human-centered AI, which emphasizes the use of AI systems to support human capabilities while maintaining meaningful human oversight (Schmager et al., 2025). From this perspective, the primary value of AI-enabled feedback is not the replacement of instructor judgment. Rather, genAI serves as a tool that can help instructors provide timely, individualized, and high-quality feedback at greater scale. Consistent with this approach, we clarified to the instructor that the intended role of the AI tool was to augment, rather than replace, their role in the feedback process and that they remained responsible for the content of the feedback provided to students.

In the present study, we use the term “AI-enabled feedback” to refer to feedback developed within a human-centered AI framework. Within this framework, instructors retain responsibility for the feedback process rather than delegating it entirely to an AI system. Their involvement may include configuring the system for a specific assignment, choosing and applying learning theory to the implementation, evaluating generated output, and revising feedback before it is provided to students. Such oversight helps ensure that feedback remains aligned with instructional goals and assessment criteria while preserving instructor accountability.

Customization of the genAI system is a particularly important aspect of this process. By supplying task-specific instructions, disciplinary resources, and contextual information, instructors can shape genAI output to reflect the needs of a particular course or assignment. Prior research suggests that these practices can improve the accuracy and relevance of generated feedback while reducing hallucinations, or outputs that sound plausible but are inaccurate (Mills et al., 2025; OpenAI et al., 2024).

#### 2.6.2 Pedagogical Alignment and Feedback Structure

The learning objectives are an important part of the context in which AI was implemented. The assignment examined in this study was guided by the following learning objectives: (1) To write a first draft of the introduction of your research proposal, and (2) To respond to the reviewer’s feedback and compile the final draft of your research proposal.

The GPT was deliberately designed to draw on course-specific materials and learning supports. M.W. worked with the course lab coordinator to develop the research proposal assignment and accompanying rubric, ensuring that both reflected the coordinator’s instructional goals. The rubric incorporated the Claim, Evidence, and Reasoning (CER) framework, which had been used throughout the course to support students’ scientific explanations (McNeill & Krajcik, 2008). To provide additional guidance, we created a student-facing scaffolding tool that described the organization of the proposal introduction and the content expected within each paragraph. The GPT was then set up to generate feedback aligned with the rubric, CER framework, and scaffolding tool. Consequently, the feedback followed the rubric’s structure, as shown in Table 3. The resulting feedback was structured so that each rubric section included: (1) identified strengths, (2) suggestions for improvement, and (3) examples intended to scaffold those improvements. The GPT was tested by reviewing the structure and accuracy of feedback generated on two sample proposals. The full instructions and configuration—including the knowledge base—is provided in the Supplemental Materials.

**Table 3.** Structure of the feedback sections provided to students.

| Feedback Section | Scope | Description |
|---|---|---|
| Macro | Paragraph-specific | Feedback pertaining to the content of the Macro paragraph. |
| Meso | Paragraph-specific | Feedback pertaining to the content of both the Meso Part 1 and Meso Part 2 paragraphs. |
| Micro | Paragraph-specific | Feedback pertaining to the content of the Micro paragraph. |
| Flow of Writing | Whole-proposal introduction | Feedback on how effectively each sentence and paragraph transitions to the next, with a clear flow of ideas. |
| Claims are Supported | Whole-proposal introduction | Feedback on how well claims are supported by scientific reasoning and evidence from credible cited sources. |

#### 2.6.3 Institutional and Course-Level AI Use Policies

Because we examined students’ experiences with and perceptions of genAI in a learning context, it is important to describe the institutional and course-level AI policies that shaped that context. At the time of the study, the university’s guidelines for genAI use did not address instructor use of AI. Instead, instructors were given discretion to establish expectations for AI use within their courses. The instructor in this study did not provide a written AI policy to students. Rather, she informed students on the first day of class that they could use genAI responsibly and trusted them to determine what responsible use entailed. She also explained that if she believed a student had violated this expectation, she would discuss the issue with the student and allow them to resubmit the assignment without penalty.

## 3. Analysis

This section begins with an overview of the themes developed from the analysis of student team interviews. We then provide an in-depth examination of each theme, followed by a synthesis of findings. Pseudonyms are used throughout to protect participant identities.

### 3.1 Themes

We developed three themes, which are presented in Table 4 and elaborated below. The themes are: (1) Students recognized characteristics of effective feedback, (2) Positive emotions encouraged intention to revise, and (3) Students accepted but critically evaluated instructor AI use.

**Table 4.** Themes constructed from student team interviews.

| Theme Name | Description |
|---|---|
| Students recognized characteristics of effective feedback | Students recognized characteristics they associated with effective feedback in the AI-enabled comments, such as comprehensiveness, detail, actionability, and personalization, which align with characteristics that prior research has identified as important to students. This suggests that students' positive perceptions of AI-enabled feedback were rooted less in the use of AI itself and more in the extent to which the feedback supported improvement. |
| Positive emotions encouraged intention to revise | The feedback fostered positive emotions about the outcome of the students’ first draft assignment and their ability to succeed in revising. Identifying the proposals’ strengths and giving actionable tips cultivated feelings of pride and confidence that helped re-engage students with the revision task. |
| Students accepted but critically evaluated instructor AI use | Students generally accepted instructor AI use but emphasized the importance of how it is implemented. They wanted the instructor to remain actively involved in the feedback process and rejected full automation. Their concerns were centered on maintaining fairness, quality feedback, and upholding the instructor’s responsibility for guiding learning. |

#### 3.1.1 Theme 1: Students recognized characteristics of effective feedback

This theme presents a pattern in students’ perceptions of the AI-enabled feedback itself. Students thought the AI-enabled feedback was more thorough because it included additional types of comments that set it apart from previous feedback they had received on their writing in other courses. Students were used to feedback that simply identified areas for improvement, but what distinguished the AI-enabled feedback, in their view, was its inclusion of strengths and emphasis on *how* to improve. For example, Linda from Team 1 said:

> *I like how I feel like most professors, when they give feedback, it’s very short and limited and they don’t really say your strengths. They just tell you what needs to be improved. So I like how we got feedback on our strengths and then also that scaffolding tip of, “Here’s your weaknesses and here’s how you can get a head start on improving it”, because most of the time they just tell you your weaknesses and then it’s like, like I said, okay, well, what do I do with this, how do I get to where I need to be?*

Related to the comprehensiveness of the feedback, students often appreciated its structured format. The structured flow—from highlighting strengths, to suggesting improvements, and concluding with scaffolding tips—formed a progression that built on itself, elaborating comments and situating them within sufficient context to guide revision. This layered structure appeared to enhance clarity. For example, Team 2 received the following feedback on the first paragraph of their proposal introduction (the bold font is from the feedback’s original formatting):

> *Strengths:…The proposal introduces redlining as a historical practice that has contributed to current health disparities, particularly diabetes. The statement that “redlining contributes significantly to the variation in diabetes mortality rates” begins to trace the connection from past to present.*

> *Suggestions for Improvement: To strengthen this section, **clearly walk the reader through the historical policies** that institutionalized redlining (such as the role of the HOLC or FHA), and **explain how these policies caused long-term economic and health disparities.** The connection between historical redlining and present-day diabetes is mentioned, but **the causal chain could be more fully explained.***

> *Scaffolding tip: You could start with, “In the 1930s, redlining was codified through federal housing policy…” then show how that led to reduced investment, fewer health services, and ultimately worse health outcomes today.*

Whereas other studies have noted that students often find feedback confusing and difficult to use (Weaver, 2006; Winstone et al., 2017), these teams consistently found the feedback easy to interpret and actionable. Mya from Team 1 explained how the feedback compared to typical comments:

> *I feel like some of the feedback I’ve received on other essays or projects tend to be more vague, but I feel like with this the way that the feedback was formatted felt that my work was valuable but also gave me exactly like what I can do to improve it.*

Similarly, Lucy from Team 2 described the value of the feedback’s structure:

> *I’ve had professors that would just kind of tell you what was wrong with it, but wouldn’t give you any help to like, “Oh, this is how you can improve it” or “This is kind of where you did good”. But I do like this because it kind of, like she said, outlines the strengths and outlines suggestions for improvement and also, kind of gives you a tip of how to actually improve it and I think that’s very helpful because getting feedback, you want to be able to yes, take the feedback and try to figure out how you can improve it, but it also helps if someone kind of gives you a little hint.*

Beyond the feedback’s overall structure, students identified two specific features as particularly supportive of revision: comments that clarified existing strengths and scaffolding tips that suggested how to move forward with revising. Students described the strengths section as helping them identify effective elements of their writing that should be kept and built upon during revision. For instance, Lucy described the influence of the strengths section on her revisions: “*I do like how it outlines the strengths like I said it shows you what you did right, maybe what you shouldn’t change about what you did.*” Acknowledging strengths can also foster positive emotional responses to feedback, a point explored further in the next theme.

While strengths clarified what was already working, scaffolding tips helped students determine how to improve areas that needed further development. In our interviews, the most frequently emphasized quality was AI-enabled feedback that went beyond identifying weaknesses to include specific suggestions for how to improve, which strongly influenced students’ perceptions of its usefulness. Reflecting a shared perception of the feedback, Linda stated:

> *I kind of like how we have that area for improvement, but then the tip kind of helps you get a start of where to go and how to make that improvement because sometimes when you’re given feedback, you kind of feel stuck and you don’t really know where to go with it. So I kind of just like how it gives you a starting point of where to go.*

Similarly, Lucy appreciated scaffolding tips, noting in one of her interview segments above that they provided “*a little hint*” that bridged the gap between receiving feedback and making improvements.

Despite his generally positive views of the feedback’s comprehensiveness, Dakota from Team 2 expressed reservations about the amount of feedback he received, particularly in comparison to his groupmates. Although the feedback for each section of the proposal followed the same format, the feedback on Dakota’s section was longer than that provided for the rest of the proposal. He explained his reaction this way:

> *I felt really kind of bummed out, but not really…I knew it was some improvement that needed to be made, but the just feedback gave me that clarification. But I did see how my feedback, it was so much longer and more detailed than my peers, as my group members and that made me feel like…it just made me think back, and look back at what I did and how much I needed to improve, so that made me feel kind of bad but it made me reflect on what I needed to improve on so that also made me feel better at the same time, if that makes sense.*

Dakota’s response suggests that comprehensiveness, especially when the amount of feedback differs across groupmates, may elicit negative feelings related to social comparison or being overwhelmed. At the same time, his experience appeared ambivalent. Although the volume of feedback initially made him feel discouraged, he also valued the clarity it provided and viewed it as helpful.

Although students generally approved of the detailed and comprehensive feedback, Dakota’s feeling of being “*bummed out*” by the amount of feedback highlights potential tradeoffs. Prior research indicates that extensive feedback may not always be effective and can sometimes leave students feeling overwhelmed (Freeman & Lathbury, 1998; Lipnevich & Smith, 2009; Lunsford, 1997). Students may also make social comparisons (Festinger, 1954), interpreting longer feedback as a signal that their work was less successful than that of their peers. In the emerging context of AI-enabled feedback, these concerns may become increasingly consequential because genAI can easily generate extensive comments. As a result, it may be important to set up AI tools to strike a balance between providing detailed, comprehensive feedback and limiting the volume of comments to a level that is manageable for students. Researchers and instructors may also need to consider how differences in feedback length across students influence students’ perceptions of their performance relative to their peers.

The patterns we observed indicate that students recognized characteristics they associated with effective feedback in the AI-enabled comments. (Dawson et al., 2019; Higgins et al., 2002; Jonsson & Panadero, 2018; Li & De Luca, 2014). Although these students did not explicitly identify personalization as important, it may have been implied in their appreciation for comments clarifying their strengths and providing scaffolding tips that suggest specific next steps. In general, the participants thought that the feedback supported improvement through its comprehensiveness, detail, actionability, and personalization, which align with characteristics that prior research has identified as important to students (Dawson et al., 2019; Ferguson, 2011) and extends it into the context of AI-enabled feedback. These findings also suggest that students’ positive perceptions of AI-enabled feedback were rooted less in the use of AI itself and more in the extent to which the feedback supported revision and improvement.

#### 3.1.2 Theme 2: Positive emotions encouraged intention to revise

In addition to supporting intention to revise through comprehensive comments, our interviews showed that AI-enabled feedback also encouraged a willingness to revise by positively influencing their academic emotions. One way the feedback fostered positive emotions was by identifying strengths in students’ first drafts. Highlighting strengths may have helped students associate positive outcomes with their own actions, contributing to feelings of pride, accomplishment, and confidence (Ferguson, 2011; Weiner, 1985). Although we previously discussed the following quote from Mya in relation to students’ interpretation of feedback, it also illustrates the positive emotional response elicited by the strengths section: “*I feel like with this the way that the feedback was formatted felt that my work was valuable*”. Later, Mya elaborated on her perception of the strengths section:

> *Mya: I thought, I personally liked how she, the feedback provided strengths.*

> *Interviewer: What do you think is helpful or good about giving your strengths?*

> *Mya: I think in a way it gives us confidence that you know we’re not, it’s not as bad as we think it is but it gave us, it gave me personally confidence and it showed me like what I needed to improve on and showed me what are the things I should keep.*

Lucy also responded positively to the strengths section of her feedback and appeared to experience both a sense of accomplishment and relief:

> *I really liked that it outlined our strengths first because I think it is I don’t know. Sometimes when you get feedback, you kind of feel like you have to change everything so it’s nice to know that there are some good parts of it or like it’s nice to know what you did right first, I guess.*

Clarifying the strengths of a first draft highlighted areas of success that may otherwise have gone unnoticed. The resulting feelings of pride and self-confidence seemed to increase students’ intention to continue improving their work (Ferguson, 2011; Hattie & Timperley, 2007; Pekrun, 2006). Pride is considered a positive activating emotion associated with interest and the use of flexible, creative learning strategies (Pekrun, 2006). In feedback contexts, these emotions may motivate students to engage with and implement feedback (Hattie & Timperley, 2007), both of which are essential for students to benefit from the feedback they receive (Boud & Molloy, 2013; Hattie & Timperley, 2007; Henderson et al., 2019).

Although the strengths section fostered positive activating emotions, it also appeared to elicit relief, a positive emotion that can have mixed effects on learning and is sometimes deactivating. In some circumstances, relief may lead to complacency and reduce students’ motivation to re-engage with the learning task (Pekrun & Linnenbrink-Garcia, 2012). During our interviews, both Mya and Lucy seemed relieved after seeing that their first drafts had been successful in at least some respects. In this case, however, the effects of relief on revision may have been limited because the strengths section focused on aspects of students’ drafts that were already successful and therefore did not require revision. Consequently, the broader influence of pride and confidence may have played a larger role in supporting students’ intention to revise.

Whereas the strengths section appeared to foster feelings such as pride, accomplishment, and confidence, the feedback’s practical support provided a separate source of confidence by helping students feel prepared to successfully address the comments. Lucy expressed a sense of clarity when she explained that a scaffolding tip “*…gives you a more of a tangible step in between the feedback and actually fixing what you did wrong*”. Similarly, Carl from Team 1 recalled, “*I just remember starting out having an idea what to do so…I was able to get to it real quickly*”. Linda also reflected on her revision experience in the following exchange:

> *Interviewer: How do you remember feeling as you use the feedback to revise the proposal? Linda: I definitely feel more confident when I was revising and rewriting so I know my weaker areas and how to improve them.*

Together, these examples suggest that students felt confident and prepared because the feedback was actionable and clearly identified what needed revision while affirming what was already working well. Control-value theory offers one explanation for these achievement emotions by linking emotions to individuals’ appraisals of control and value within a learning context (Pekrun, 2006). In this context, the feedback’s practical support may have enhanced students’ perceived control over successful revision, thereby motivating them to engage in the revision process.

Although students discussed positive emotions and experiences more frequently than negative ones, they also described some negative feelings. One pattern involved frustration when feedback requested information that students had already researched but were unable to find. For example, the feedback suggested that Team 2 find a personal story from a local resident, rather than a resident of a different city, to illustrate the health outcome and ecological impact at an individual level. It went on to say: “You can paraphrase local interviews, use testimonies from food bank users or gardeners, or reference community members quoted in local news articles.” This feedback pertained to the paragraph of the proposal authored by Rhyan from Team 2. During the interview she explained:

> *I was kind of a little irritated because I knew I was trying to, I already had tried to find it and everything so when I had seen it again, to try and find an individual action in [city], I was like, I’ve been trying to do that so it’s, I don’t know.*

Lucy described a similar experience after receiving feedback suggesting that she more thoroughly describe the community response initiative in her paragraph. She found the feedback frustrating because, as she explained, “*…I’ve already tried to look for examples of this or policies or whatever. And there’s just no information about it*.*”*

Although frustration is a negative emotion, it can be activating, prompting students to direct effort back to the task (Pekrun & Linnenbrink-Garcia, 2012). Despite potentially low perceived control over locating the requested information, both Lucy and Rhyan reported conducting additional research to try to address the feedback. As Rhyan explained:

> *But I did go back, I continued to try to type in different ways into the search bar like try to see if I can find something I missed when I was doing my research or just trying to find a different article maybe, but I still didn’t find it.*

In this context, even a negative emotional response to feedback requesting hard-to-find information appeared to encourage continued engagement with the revision task.

#### 3.1.3 Theme 3: Students accepted but critically evaluated instructor AI use

When reflecting on instructor use of genAI after participating in this project, these students tended to be more concerned with how tasks and responsibilities were distributed across genAI and their instructor than merely AI’s presence in the process. Students articulated their own expectations regarding acceptable and unacceptable instructor uses of AI and evaluated the feedback process accordingly. For instance, Dakota said:

> *So for feedback, I feel like it’s okay to use [genAI] for certain things for feedback. But you said that Dr. Susie did, our professor did revise or read or look at our work before putting it into AI, so, I feel like that was fine. And then if that wasn’t the case, and it was just a copy and paste type of thing then I still don’t agree with that, using AI for that. Just because professors are supposed to teach us what to do, not AI.*

In this example, Dakota rejected full automation—a view that was common among participants. Specifically, students wanted their instructor to read each proposal and remain actively involved in the feedback process. Their concerns centered on maintaining feedback quality and ensuring that their instructor remained responsible for guiding student learning. The excerpt below from Mya also describes an expectation that the instructor is ultimately responsible for providing quality feedback:

> *If it’s [AI-enabled feedback] something that they look into again and they see how if it’s relevant or not or if it’s anything that they can add, then I think it’s a good tool. But if it’s like if you just putting it in there and then copy and pasting the feedback, then I don’t think it’s a helpful tool for instructors cause when I think of an instructor, I think of someone who has like an education behind what they’re doing, like whether that’s a PhD or like a degree. So like when I see someone with that, I would think that they’d be able to give good feedback by themselves. But if they’re using it, how this was used, I don’t mind.*

Another area of concern for students was the responsibility of assigning grades. Although this project involved formative feedback and grades were not at stake, students consistently emphasized that they did not want genAI to assign their grades. Linda said it this way, “*As long as it’s not giving me grades on things, I’m fine with it. It’s not sitting there being like, this is their work, this is the grade you should give them.*”

Dakota expressed an additional criterion for evaluating instructor AI use that is noteworthy. He considered instructor use of genAI in relation to the course’s expectations for student genAI use. In particular, he raised concerns about the fairness of allowing instructors to use genAI while restricting its use by students. Dakota explained:

> *If they don’t allow us to use AI, I don’t think they should use AI to grade us…I think they could use it. I think I’m not opposed to using it, like having some use, but not completely. Same thing like with us like paraphrasing things like that. So I think it applies to both sides…and complete use meaning as not even looking over our work, copy and pasting.*

Although most participants articulated expectations regarding how tasks and responsibilities should be distributed between genAI and their instructor, there was variation in how they evaluated instructor AI use. In particular, Carl stated that he had few concerns or preferences, aside from trusting a custom GPT, like the one used in this study, more than standard ChatGPT, which is not designed for a specialized task. When explaining his seemingly unreserved acceptance of instructor use of genAI, he stated:

> *I mean I’m used to like teachers just scraping random worksheets off the internet to give to their students as homework assignments so having ChatGPT give feedback or make an assignment, it’s not too much different. As long as it’s not judging my grade, I don’t care.*

Carl also compared instructor AI use to the assistance they receive from course ULAs (Undergraduate Learning Assistants). He explained, “*I genuinely assume that the ULAs do a lot of the work so ChatGPT, ULA, it doesn’t feel too much different.*” While Carl’s comments suggest that some students may not hold strong preferences regarding acceptable instructor uses of AI, they may also indicate that he did not perceive this particular implementation as violating any important instructional responsibilities.

These students generally expressed acceptance of instructor AI use, and none categorically rejected it. Their acceptance appeared to be shaped by two factors: the quality of the AI-enabled feedback they received during the project, and their prior experience using genAI. While the role of students’ experiences with this AI-enabled feedback is explored further in the Synthesis section, students’ prior use of genAI also appeared to influence their acceptance of instructor AI use. Previous experience seemed to normalize the technology, and they often described it as well suited for writing-related tasks. In fact, two students discussed using genAI to provide feedback on their own writing. Rhyan explained her perspective this way:

> *In times for feedback like this, I think that it’s okay to use [genAI] because it’s kind of the same as if I were using and putting my paragraph in there to get feedback for changes and stuff so I can, I like the fact that though they [the instructor] read it first instead of putting it in the AI and then, giving feedback and taking the feedback from there. But I think that it’s kind of, it can be very useful.*

Prior research has shown that students’ perceptions of a feedback provider, particularly the provider’s credibility, influence perceived feedback effectiveness (Telio et al., 2016; Watling et al., 2012). However, little is known about how instructor use of genAI shapes these perceptions. As a result, the implications of genAI for the feedback process remain unclear. Our findings help address this gap by showing that, in the context of AI-enabled feedback on scientific writing, students considered both the instructor’s credibility and the credibility of genAI. In addition, instructor oversight was a key factor in their evaluations. Participants generally assumed that their instructor was credible and capable of providing quality feedback. This view was exemplified by Mya’s comment that “*when I think of an instructor, I think of someone who has like an education behind what they’re doing…I would think that they’d be able to give good feedback by themselves.*” In contrast, students expressed mixed views about genAI’s credibility as a feedback provider. Although they typically viewed genAI as well-suited for writing-related tasks, most did not trust the feedback it generated without oversight. Consequently, students wanted instructors to read their proposals and vet the feedback generated by genAI. These findings suggest that credibility remains an important consideration in AI-enabled feedback contexts but becomes more complex when genAI is introduced into the process. Students may therefore evaluate the overall feedback process not only in terms of the instructor’s expertise, but also in terms of whether the instructor’s involvement addresses perceived limitations of genAI.

### 3.2 Synthesis of Themes

Collectively, students evaluated AI-enabled feedback according to whether it supported revision and instructor responsibility. This synthesis is illustrated in Figure 1, which shows the interconnectedness of the three themes that shaped students’ overall experiences with AI-enabled feedback. The left panel presents the two themes that pertain to students’ experiences with and perceptions of the feedback itself, and the explanatory relationship between them. Students recognized characteristics of effective feedback in the AI-enabled comments, which made it seem useful for revision. This practical support may have enhanced students’ perceived control over successful revision, often eliciting positive emotional responses that motivated students to use the feedback (Pekrun, 2006). Together, the themes suggest that AI-enabled feedback supported intention to revise through both its content and its effect on students’ academic emotions.

**Figure 1.**
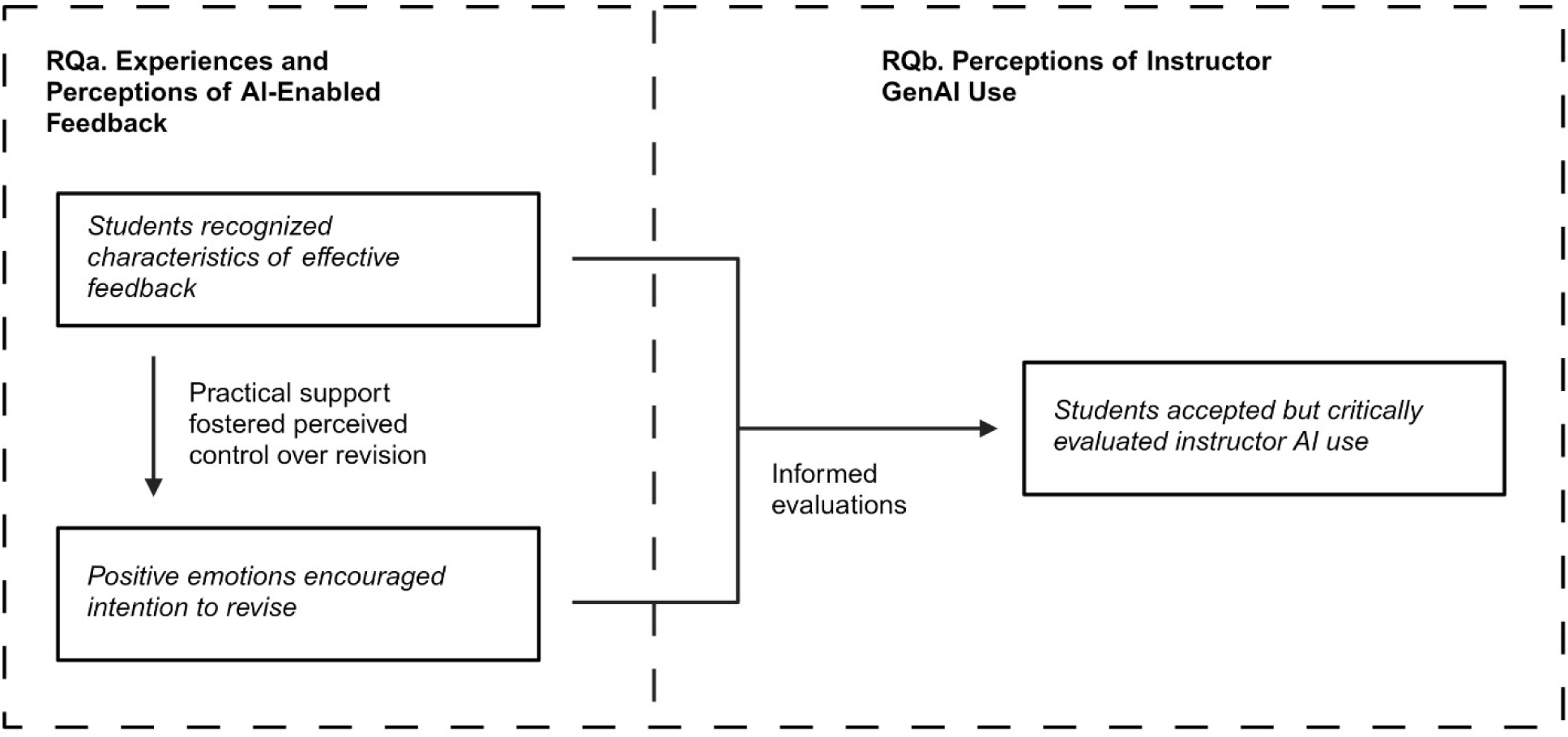
Themes developed from student team interviews. Themes are shown in boxes. Italicized text indicates theme names. Arrows indicate relationships between themes.

Because our evidence for students’ feelings of preparedness and confidence to engage in revision came from retrospective interviews, participants’ recollections of the revision process were especially valuable. For example, Linda shared, “*I definitely feel more confident when I was revising and rewriting,*” while Carl remarked, “*I was able to get to it real quickly.*” Findings reported in Weston & Long (2026), which examined revisions made to the same students’ proposal introductions in this project, provide additional support for this interpretation. In that study, we found that both student teams made substantive revisions—those that altered the meaning of the writing—to their proposal introductions and addressed approximately half of the feedback comments. Together, these investigations suggest that intention to revise—shaped by the AI-enabled feedback—was accompanied by behavioral engagement on the revision task. This combination of intention to revise and behavioral engagement provides insight into how AI-enabled feedback can support students’ scientific writing, particularly through feedback that students perceive to be effective and emotionally supportive.

Although this study examined AI-enabled feedback, the qualities students appreciated and the affective responses they described aligned with desired characteristics of feedback more broadly. Yet a persistent challenge remains: although researchers have identified student beliefs about characteristics that make feedback effective, students often continue to receive comments that do not support their efforts to improve, and sometimes they do not receive timely feedback (Carless, 2006; Derham et al., 2022). These gaps may partly reflect practical constraints instructors face when providing feedback. In particular, instructors frequently lack the time needed to develop detailed, personalized comments (Hanmore et al., 2024). Although this study did not aim to compare the time required to produce AI-enabled feedback versus traditional feedback, and our analysis is limited to two student teams, the approach has the potential to scale personalized formative feedback. This structure ensured that both teams, regardless of how closely their work met the goals of the assignment, received comments that they found effective. Our findings suggest that, when grounded in course-specific materials and support, genAI has the potential to support instructors in developing feedback that students perceive as useful and motivating for improving their scientific writing.

Because students largely evaluated the feedback positively, they were also more willing to view their instructor’s use of genAI favorably. The right panel of Figure 1 presents the theme related to students’ perception of their instructor’s AI use, and its connection to their evaluation and experiences of the feedback itself. Students’ positive perception of the feedback’s content and their affective responses seem to have contributed to their acceptance of their instructor’s use of genAI. The detailed and actionable nature of the feedback may have helped convince students that genAI could be a useful tool for supporting instructor feedback. This pattern further suggests that students’ acceptance of instructor AI use may be shaped by their experiences with the feedback itself, rather than by the use of AI alone. Lucy described how her experience with the feedback influenced her perspective:

> *And I feel like the feedback that we got for our sections like our initial feedback, it was pretty helpful and it was pretty easy to implement so if that [genAI] was something that helped create that feedback, then I’m like, I think it was like beneficial.*

Similarly, Linda said:

> *I like the feedback that was given through the, the use of the AI that she used, so I think if it’s used in a way like this where it gives you your strengths, improvement and then tips, I really like that, and I feel like ChatGPT just is really…strong in writing areas like this, gives more information, and more like detailed descriptions.*

When considered together, these three themes extend a learner-centered perspective on feedback to contexts in which genAI is involved in developing feedback. Such an approach would emphasize not only the feedback itself but also students’ perceptions of and engagement with it (Boud & Molloy, 2013). Our findings suggest that students are more likely to respond positively to AI-enabled feedback when they perceive it as supporting improvement and aligning with their expectations regarding responsibility, quality, and fairness. Taken together, these themes suggest that students’ experiences and perceptions of AI-enabled feedback provide important insight into how genAI can be incorporated into feedback processes in ways that are both acceptable to students and supportive of their intention to revise their writing.

### 3.3 Future Research

GenAI is increasingly being used to develop feedback and can generate large amounts of personalized output in a short time (e.g., Kasneci et al., 2023; Mills et al., 2025; Wan & Chen, 2024; Xia et al., 2026). However, our interviews suggest that more feedback is not always better. Students may feel overwhelmed by extensive comments and discouraged when they compare the amount of feedback they receive with that of their peers (Festinger, 1954; Freeman & Lathbury, 1998; Lipnevich & Smith, 2009; Lunsford, 1997). As genAI enables instructors to provide comprehensive feedback at scale, it will become increasingly important to understand the processes through which students engage with, interpret, and act on feedback. This understanding will be foundational for integrating genAI into feedback processes in ways that are most effective for supporting improvement.

We found that students’ perceptions of instructor genAI use were influenced by the extent to which its implementation aligned with their expectations for fairness and responsible AI use, particularly the expectation that instructors remain actively involved in the feedback process. In addition, students’ evaluations of the feedback sender’s credibility reflected their perceptions of both the instructor and the genAI tool, including the nature of the instructor’s interaction with the tool. These findings raise important questions about how varying levels of alignment with students’ expectations shape their perceptions of both the feedback and its source. They also prompt questions about how those perceptions, in turn, influence students’ intention to engage with feedback and its effectiveness in supporting revision. Addressing these questions would provide insight into how a broader range of perceptions of genAI implementations affect students’ acceptance and uptake of feedback.

The present study relied on interviews rather than revision artifacts and focused on students’ motivation to revise. In a prior analysis of the revisions produced by the same students participating in this study, we found that students frequently acted on the AI-enabled feedback and made substantive changes to their writing (Weston & Long, 2026). While these studies together provide insight into how students’ experiences and perceptions of the AI-enabled feedback may have shaped their revisions, the interviews were retrospective. This means we were unable to directly observe how students used to feedback while revising. Future work could investigate the mechanisms of how students use feedback to revise their writing, which would inform how to better design AI tools to support this process.

## 4. Conclusion

Taken together, the themes developed in this study suggest that students’ experiences with AI-enabled feedback are shaped by its content and its influence on their academic emotions, both of which supported their intention to revise their proposal introduction. Looking back on their experiences, students considered the credibility of both the instructor and the genAI tool, viewing instructor oversight as a crucial factor. Ultimately, students’ positive perceptions of the AI-enabled feedback and its use by the instructor were rooted less in the mere presence of AI and more in their perception of its quality and support of improvement. In this context, when students perceived AI-enabled feedback as useful and its implementation as aligned with their expectations for appropriate and responsible AI use, they were inclined to engage with it—a necessary component of a learner-centered feedback approach. Collectively, these findings support the value of a learner-centered approach to genAI implementation in the feedback process, one that actively attends to students’ expectations and perspectives as feedback recipients.

## Author Notes

We have no conflict of interest to disclose.

This material is based upon work supported by the National Science Foundation Graduate Research Fellowship Program under Grant 2235783, the SimBio Foundation grant program, and the Plant Biology Department of Michigan State University.

## Supporting information

Weston and Long Supplemental Materials

## Acknowledgements

**Use of AI in the writing process:** During the preparation of this work, we used Copilot Chat (basic) to improve the clarity and conciseness of the writing and to provide critiques of the clarity of arguments. Both authors reviewed AI suggestions and edited accordingly. We take full responsibility for the content of the publication.

