## Supplementary material for "Actionable, Motivating, and Not Beyond Question: Student Insights On AI-Enabled Feedback and Instructor AI Use": Weston and Long Supplemental Materials

### Feedback-Generating GPT

*ChatGPT* (Version GPT-4o). (2025). [Large Language Model]. OpenAI.

**Contents:**


1. IRB Approval and Protecting Student Data
2. System Configuration
3. Overview of the Steps, Prompting Techniques, and Testing I Used to Refine the GPT's Instructions
4. List of Files in the GPT's Knowledge Base
5. Final Version of the Instructions


1. IRB Approval and Protecting Student Data

This proposed work was approved and found exempt by Michigan State University's Institutional Review Board (STUDY00010636). The use of generative AI in this study was approved by MSU's IT Governance, Risk, and Compliance office. We protected student data by only analyzing proposals for which we obtained written consent from students for the use of genAI in our research, by de-identifying the data prior to its analysis with GPT, and by excluding the chat conversations from OpenAI's training data.

2. Overview of the Steps, Prompting Techniques, and Testing I Used to Refine the GPT's Instructions.
   - 2.1. These are two resources I used to inform my prompting techniques:
     - 2.1.1. *Brownell, B. (n.d.). 5 Advanced Prompts for ChatGPT to Boost Answers in 2025. Descript. Retrieved May 5, 2025, from https://www.descript.com/blog/article/5-advanced-prompts-to-get-better-answers-from-chatgpt*
     - 2.1.2. *Liu, D. (2023, April 27). Prompt engineering for educators – making generative AI work for you – Teaching@Sydney. Teaching@Sydney. https://educational-innovation.sydney.edu.au/teaching@sydney/prompt-engineering-for-educators-making-generative-ai-work-for-you/*
   - 2.2. Steps of developing the instructions:
     - 2.2.1. Write the initial instructions (what is your role?, what exactly do I want you to do?, what format should your output be in?)
     - 2.2.2. Revise the instructions to take on the **Role, Task, Requirements, and Instructions Format** (Liu, 2023) and embed the **Reflection Pattern** within it in the Instructions section (Brownell, n.d.). The Reflection Pattern asks the model to explain its reasoning, which helps you understand and evaluate its output.
     - 2.2.3. Improve my instructions using the **Question Refinement Pattern** (Brownell, n.d.), which entails giving the model access to information about the assignment and asking it to improve my instructions to include information relevant to the assignment. I gave it access to the feedback rubric and proposal writing scaffolding tool, which is written specifically for this project. I asked the model to make my instructions better by including information specific to this project/assignment. I used the model's output as my new version of the instructions.

2.2.4. I used the **Reflection Pattern** again, but this time as a separate prompt, not as something embedded in the instructions themselves. I asked the GPT, "When you do your job, how do you go about using your instructions?". The output was a detailed step-by-step description of the process it follows, which was helpful for shedding light on what happens "behind the scenes".

2.2.5. I also used the **Flipped Interaction Pattern** (Brownell, n.d.) to have the model ask me questions that will help it better understand its task. I updated my instructions based on what I learned from the questions the model asked me.

2.2.6. During this step I reached the 8,000-character limit for instructions. I had to make strategic decisions about what to cut and how to save space. About half of the suggestions that came out of this step were incorporated into existing sections of the instructions, while others were tacked on at the end.

2.2.7. I tested the GPT by asking it to provide feedback on an example proposal it had not seen before (that I used a different GPT to help write).

3. System Configuration

This section shows the settings under the "Configure" tab of the GPT's menu.

3.1 Conversation starters: none

3.2 Capabilities

3.2.1 Checked options: Web Search and Canvas

3.2.2 Unchecked options: 4o Image Generation and Code Interpreter & Data Analysis

3.2.3 Actions: none

4. List of Files in the GPT's Knowledge Base

4.1 Assignment artifacts:

4.1.1 Scoring rubric with category descriptions

4.1.2 Student-facing scaffolding assignment for help with writing. This assignment specified what information belongs in each paragraph.

4.2 Example proposal and its corresponding feedback

4.3 Book chapter: "Inquiry and Scientific Explanations: Helping Students Use Evidence and Reasoning" (McNeill and Krajcik, 2007)

4.4 Journal article that students read related to the topic: "The ecological and evolutionary consequences of systemic racism in urban environments" (Schell et al. 2020)

4.5 Open-source textbooks:

4.5.1 Environmental Biology by Matthew R. Fisher

4.5.2 Biology 2e by Clark, Choi, and Douglas (spring semester 2025 only)

4.5.3 Microbiology by Parker, Schneegurt, Thi Tu, Lister, and Forster (spring semester 2025 only)

5. Final Version of the Instructions

This assistant provides a draft of student-facing feedback on the Introduction section of lab proposals.

Role: Act as an expert assistant to a college biology laboratory instructor, with a focus on providing detailed feedback to students on their research proposals. Feedback must be aligned with the SS25 Intro Rubric.pdf and must also incorporate scaffolding techniques described in the ScaffoldingToolGPTversion.pdf to support students in meeting assignment expectations.

Goals:
1. Detailed Feedback Using Rubric Criteria: For each proposal, provide feedback on the entire document using all eight criteria listed in SS25 Intro Rubric.pdf ("Macro scale: Past to present", "Macro scale: Location-specific", "Meso scale: Local policies and responses", "Micro scale: Individual experiences", "Research Questions", "Variables", "Flow of writing", "Claims are supported") with a clear explanation for the score given (Complete, Needs some work, Not addressed). Use the rubric's macro, meso, and micro scale approach for evaluating content, while equally addressing each level and the connections across levels. Ensure students understand where they fall short and how they can improve.

2. Scaffolding for Proposal Improvement: Integrate the ScaffoldingToolGPTversion.pdf approach into the feedback. The tool provides a step-by-step method to help students understand complex ideas (e.g., how to structure the introduction). Utilize these steps, especially for multi-scale analysis (macro, meso, micro), to guide students in constructing well-organized arguments. Also incorporate the claim, evidence, and reasoning framework (described in the pdf in the GPT's knowledge called "Claim, Evidence, Reasoning Framework.pdf") to help students write complete scientific claims that are supported with evidence and explained via reasoning with scientific principles. But remember, students don't know the name of this framework. Describe it qualitatively to them, not by name.

Feedback Structure:
1. Score each of the eight criterion in the rubric (ex. "Macro scale: Past to present"): Use one of three scores—Complete, Needs some work, or Not addressed.
2. In your explanation, include:
 a. Strengths: Highlight what the student did well, providing specific examples to reinforce positive aspects.
 b. Suggestions for Improvement: Point out gaps or weaknesses in a supportive way. Use the claim, evidence, and reasoning framework (described in the pdf in the GPT's knowledge called "Claim, Evidence, Reasoning Framework.pdf") and the ScaffoldingToolGPTversion.pdf to provide suggestions on how to improve. For example, if a student hasn't connected macro-level policies to their local context, suggest adding specific examples from their region.
3. Give examples to bolster your suggested improvements.
4. Use SS25 Example Feedback.pdf as an example of the structure of your feedback.

Specific Guidelines to Incorporate:
1. Macro, Meso, and Micro Scales in Feedback:
  a. Ensure the introduction of each proposal covers macro-scale issues (historical/societal context) and their evolution into present-day inequities. Use scaffolding techniques to suggest how students can deepen their explanation if it is missing or needs work.
  b. The meso scale should include local policies or community responses. If missing, suggest specific ways they can integrate more local details.
 c. For the micro scale, focus on individual experiences or data, providing specific guidance on how to make personal impacts clear.
2. Provide guidance on supporting claims with evidence and reasoning
  a. When students make a claim, help them to support the claim with evidence (data from their own investigation or by citing someone else's investigation) and reasoning that "links the claim and evidence

and shows why the data count as evidence to support the claim. Often in order to make this link, students must apply appropriate scientific principles" (McNeill & Krajcik, 2008)
3. Use the file Example Proposal.pdf and its corresponding feedback in the file SS25 Example Feedback.pdf to guide the structure and content of your feedback.

Here are a few directions for when and how to query the source files in your knowledge base for relevant information:
1. Science discipline-specific content: If the proposal contains science discipline-specific content (biology, environmental biology, or microbiology) that you need to assess for your feedback, consider querying one of the attached textbook files (Biology Textbook.pdf, Environmental Biology Textbook.pdf, Microbiology Textbook.pdf) for help accordingly.

2. Topic-specific content: These independent projects fall within the broad topic of the ecological and human health consequences of systemic racism in urban environments. When you have the need to incorporate this topic into your feedback, please query the document called "Ecology, human health, and racism.pdf".

Here's more clarification on how to approach your task:
1. Intersection of Race, Environmental, and Health Outcomes: You should maintain a focus on how the proposals explore this intersection, paying close attention to how systemic racism, environmental conditions, and human health outcomes are addressed across macro, meso, and micro levels.

2. Detailed and High-Level Feedback: You will give higher-level feedback on whether the proposal meets key criteria in the rubric and also provide detailed, step-by-step scaffolding on how to improve sections, such as how to more effectively explain claims, support them with evidence, and show reasoning behind connections.

3. Content and Structure: Focus on Rubric Criteria and Proposal Support of Claims:
a. Rubric Focus: The main focus will be on how well the proposal meets the content expectations outlined in the rubric (such as explaining the macro, meso, and micro scales, identifying research questions, describing variables, etc.). You will ensure that each criterion from the rubric is evaluated, and you will provide detailed guidance on any areas that need improvement.
b. Supporting Claims: You will evaluate the support for claims, focusing on how well the proposal uses evidence and reasoning to justify its claims, ensuring that they are clear, well-supported, and linked to the research questions.

4. Item Source Usage:
a. SS25 Intro Rubric.pdf: Use this file to guide the structure of your feedback. Include all 8 criteria exactly as they appear in the rubric (Ex. "Macro scale: Past to present", etc.)
b. ScaffoldingToolGPTversion.pdf: This will guide your feedback on content and structure—ensuring the proposal effectively addresses macro, meso, and micro levels, and helping students understand how to improve.

c. Claim, Evidence, Reasoning Framework.pdf: You will use this source specifically to guide feedback on students' use of claims, evidence, and reasoning, helping them build strong, logically sound arguments in their proposals.

5. Querying the web: Please incorporate information that is specific to the project covered in the proposal you are giving feedback on, such as information specific to the location/context of the proposal.

### Proposal Drafts with Revision Coding

Brackets, such as those used in [city] and [citation #1], indicate an edit made to the draft by the researcher to obscure the location of the study.

#### Coding Legend

- **Bold** indicates addition (including meaning changes and citation changes)
- ~~Strikethrough~~ indicates deletion (including meaning changes and citation changes)
- <u>Underline</u> indicates substitution
- *Italics* indicates move

#### Team 1 First Draft

#### Macro Paragraph

~~At the macro level there is more redlining and these cities are being subdivided into regions that are being classified based on the residents ability to pay back their loans. These residents in red areas are no longer able to get these loans. “When lenders redline, mortgage loans with reasonable terms often are not available for any house in a neighborhood, regardless of the creditworthiness of the borrower or condition of the building.”~~ This is a discriminatory lending policy whereby financial organizations deny people mortgage loans or other financial services because of where they live, not because of their creditworthiness or the state of the property. Redlining disproportionately affects minority communities and low-income areas, reinforcing racial and economic segregation.

#### Meso Part 1 Paragraph

The construction of [local highway] through [city]’s historically redlined African-American neighborhoods is an example of a policy that disrupted local resources, community health, and environmental conditions. ~~Between 1963 and 1967~~, the project forcibly displaced 890 homes, demolished businesses and churches, and dismantled a once-thriving, ~~tightly-knit~~ community ~~([citation #1])~~. This loss of local resources weakened social networks, limited economic opportunities, and reduced access to culturally significant services. The displacement, coupled with inadequate relocation assistance, pushed many residents into unstable housing situations, increasing stress and economic hardship. Health outcomes worsened due to increased air pollution from highway traffic, contributing to higher asthma rates in an already vulnerable population ~~([citation #1]).~~ The area's historically lower tree density—~~resulting from redlining and disinvestment —~~also left residents exposed to more UV radiation and urban heat. These environmental challenges, ~~combined with economic barriers~~, made it difficult for residents to mitigate the harmful effects of <u>pollution</u> and extreme heat. The legacy of [highway]'s construction continues to shape disparities in resources, health, and environmental conditions, echoing the persistent effects of discriminatory policies in [city].

**Meso Part 2 Paragraph**
The community has actively responded to the negative impacts of the [highway] urban renewal project and the broader effects of redlining through initiatives like the [local garden program]. By supporting over 100 gardens across more than four acres of vacant lots in [city], this program fosters community collaboration and revitalizes neglected spaces ~~([citation #2])~~. ~~These gardens range from small household plots to larger urban farms,~~ offering residents a way to reconnect with their community, strengthen social ties, and improve local resources ~~([citation #2])~~. ~~The program also provides practical resources like soil tests, compost, and plant starts, empowering residents to actively shape their environment ([citation #2]). This initiative not only helps address food insecurity and beautifies neighborhoods but also counters the environmental degradation left by disinvestment and displacement. By reclaiming vacant land for productive use, the community works toward healing from past trauma and rebuilding the strong, interconnected neighborhoods that once characterized the area before the highway's construction.~~

**Team 1 Final Draft**
Macro Paragraph
Redlining is a discriminatory lending policy that denies people mortgage loans or other financial services based on their location, rather than their creditworthiness or property condition. This disproportionately affects minority communities and low-income areas, reinforcing racial and economic segregation. *The construction of [highway] through [city]'s historically redlined African-American neighborhoods disrupted local resources, community health, and environmental conditions, forcibly displacing 890 homes, demolishing businesses and churches, and dismantling a once-thriving community. This loss of local resources weakened social networks,*
*limited economic opportunities, and reduced access to culturally significant services.*

Meso Part 1 Paragraph
The displacement, coupled with inadequate relocation assistance, pushed many residents into unstable housing situations, increasing stress and economic hardship. Health outcomes worsened due to increased air pollution from highway traffic, contributing to higher asthma rates in an already vulnerable population. The area's historically lower tree density also let residents exposed to more UV radiation and urban heat, making it difficult for residents to mitigate the harmful effects of <u>pollen</u> and extreme heat.

Meso Part 2 Paragraph
*The legacy of [highway]'s construction continues to shape disparities in resources, health, and environmental conditions, echoing the persistent effects of discriminatory policies in [city].* The community has actively responded to the negative impacts of the [highway construction project] and the broader effects of redlining through initiatives like the [local garden program]. By supporting over 100 gardens across more than four acres of vacant lots in [city], this program fosters community collaboration and revitalizes neglected spaces, offering residents a way to reconnect with their community, strengthen social ties, and improve local resources

**Transition**
**Although there are collective efforts like the [local garden program] that demonstrate community resilience, there are still lingering effects on individuals' experiences of displacement and ecological harm.**

Micro Paragraph
In 2013, a nasty storm wiped out power for most of [city] for days on end due to improperly trimmed tree branches falling onto homes and power lines. The company had hired a general contractor to trim the trees instead of a professional landscaper, causing controversy and highlighting the need for more responsible tree trimming practices.

**Team 2 First Draft**
Macro: Redlining in the past has had a long-lasting impact on health inequities, especially the prevalence of diabetes in underserved neighborhoods. ~~According to research, redlining contributes significantly to the variation in diabetes mortality rates, and human capital is impacted by limited economic mobility (Egede et al., 2023).~~ African Americans in [city] were restricted to particular neighborhoods due to discriminatory housing practices, which limited their possibilities ~~([citation #1])~~. Diabetes disparities are sustained by ~~structural racism through~~ systemic elements like as healthcare access and poverty ~~(Ogungbe et al., 2024). According to [citation #2]~~, historically redlined regions in [city] have <u>higher incidence of diabetes</u>.

Meso part 1: In the greater [city] area, hunger is a large problem. Historical redlining practices have caused some areas to be more impacted by this issue than others. These previous practices have had effects such as less grocery or food stores available in these areas, and even potentially worse soil conditions. Soil conditions can affect the quantity and quality of food produced and worms are an organism that can significantly improve the quality of soil. If the previously redlined areas or zones that appear red on the map have worse quality soil than the other areas, food and hunger could be a problem. While this might not sound like an issue that would lead to obesity, it can cause people to turn to less healthy and nutritious food options from food banks or grocery stores.

Meso Part 2: To combat the hunger problems in the greater [city] area, ~~many~~ community gardens have been added. Some of these community gardens donate produce to the [local food bank]. These gardens provide a positive community space as well as producing food. Although many gardens have been added, they tend to be spaced away from the red-zoned areas. Gardens in the red-zoned areas tend to be private, leaving these areas with fewer gardens to contribute to the food banks.

Micro: Redlining has had an effect on food access and quality in many minority communities across the nation. The lack of good quality food can be linked to health disparities, such as diabetes and high blood pressure, since it is hard to find nutritious food in these communities. Stephanie Frias talks about this in her article about How Redlining Negatively Impacts Food Environments for Minorities. She directly states that " Growing up accessing nutritious food was
sometimes a hefty obstacle for my family"( Frias, 2024). This is showing us how finding healthy food isn't always accessible for others. In order for her to find healthy food she states that " For me I would have to take a bus, a train, and then another train to get to my nearest Whole Foods, which is about a 40-minute commute on New York City transit" ( Frias, 2024). Which is showing that people have to go out of their way to just have access to nutritious food.

**Team 2 Final Draft**

Macro: **In the 1930s, the Home Owners' Loan Corporation (HOLC) and the Federal Housing Administration (FHA) institutionalized segregation through redlining, classifying neighborhoods into four color-coded zones. Red and yellow areas, home to low-income Black residents, faced mortgage discrimination, while blue and green zones housed wealthier white populations.** This segregation has had lasting effects, contributing to health inequities like **higher obesity** and diabetes rates due to systemic barriers such as **limited** healthcare access and economic hardship. In **Greater** [city], redlined communities have significantly <u>higher obesity rates</u> than green zones, **highlighting the ongoing impact of discriminatory housing practices.**

Meso part one: In the greater [city] area, hunger is a large problem. Historical redlining practices have caused some areas to be more impacted by this issue than others. These previous practices have had lasting effects on things such as access to grocery stores and soil conditions in previously red lined areas. Soil conditions can affect the quantity and quality of food produced and worms are an organism that can significantly improve the quality of soil. If the previously redlined areas or zones that appear red on the map have worse quality soil than the other areas, food and hunger could be a problem. While this might not sound like an issue that would lead to obesity, it can cause people to turn to less healthy and nutritious food options from food banks or grocery stores.

Meso part two: In response to the issues with hunger, the [city] community implemented community gardens that donate to the [local food bank], **which aims to provide food pantries in the area with fresh produce. The [local food bank] is a non-profit organization that works with the community to provide food to those in need.** Through these community connections, the development of gardens that donate to the food bank began. These gardens provide a positive community space as well as producing food. Although many gardens have been added, they tend to be spaced away from the red-zoned areas. Gardens in the red-zoned areas tend to be private, leaving these areas with fewer gardens and less produce.

Micro: Redlining has had an effect on food access and quality in many minority communities across the nation. The lack of good quality food can be linked to health disparities, such as diabetes and high blood pressure, since it is hard to find nutritious food in these communities. Stephanie Frias talks about this in her article about *How Redlining Negatively Impacts Food Environments for Minorities.* She directly states that " Growing up accessing nutritious food was sometimes a hefty obstacle for my family"( Frias, 2024). This is showing us how finding healthy food isn't always accessible for others. In order for her to find healthy food she states that " For me I would have to take a bus, a train, and then another train to get to my nearest Whole Foods, which is about a 40- minute commute on New York City transit" ( Frias, 2024). Which is showing that people have to go out of their way to just have access to nutritious food. **Even though I wasn't able to get a personal statement from someone in the greater [city] area, these conditions and external factors also were talked about in many articles that talked about the redlining impacts on the [city] area and midwestern area.**
